# Investigation of the neural effects of memory training to reduce false memories in older adults: Univariate and multivariate analyses

**DOI:** 10.1101/2022.11.08.515495

**Authors:** Indira C. Turney, Jordan D. Chamberlain, Jonathan G. Hakun, Ashley C. Steinkrauss, Lesley A. Ross, Brenda A. Kirchhoff, Nancy A. Dennis

**Affiliations:** Department of Neurology, Columbia University, New York, NY, 10032; Department of Psychology, The Pennsylvania State University, University Park, PA, 16802; Department of Neurology, The Pennsylvania State University, Hershey, PA, 17033; Department of Psychology, Clemson University, Clemson, SC, 29634; Department of Psychology, Saint Louis University, St. Louis, MO, 63108

**Author notes:** Corresponding Author: Nancy A Dennis Department of Psychology, The Pennsylvania State University 450 Moore Building, University Park, Pennsylvania 16802.

**Keywords:** false memories, cognitive training, monitoring, univariate, multivariate, aging

## Abstract

The growing population of older adults emphasizes the need to develop interventions that prevent or delay some of the cognitive decline that accompanies aging. In particular, as memory impairment is the foremost cognitive deficit affecting older adults, it is vital to develop interventions that improve memory function. This study addressed the problem of false memories in aging by training older adults to use details of past events during memory retrieval to distinguish targets from related lures. We examined the neural basis of a retrieval-based monitoring strategy by assessing changes in univariate BOLD activity and discriminability of targets and lures pre and post training. Results showed training-related decreases in false memory rates with no alterations to hit rates. Training and practice were associated with altered recruitment of a frontoparietal monitoring network as well as benefits to neural discriminability within network regions. Participants with lower baseline neural discriminability between target and lure items exhibited the largest changes in neural discriminability. Collectively, our results highlight the benefits of training for reductions of false memories in aging. They also provide an understanding of the neural mechanisms that support these reductions.

## 1.0 Introduction

Memory impairment is the foremost cognitive deficit affecting older adults (Jacoby & Rhodes, 2006), with research showing that age-related memory impairment arises equally from age-related increases in forgetting and increases in false memories (McCabe et al., 2009). A false memory is a memory for something that never actually happened. Examples include remembering that you left your keys on the kitchen counter when you left them on the dining table, or remembering that the doctor said to take your pills in the evening when in fact she said to take them in the morning. In most cases, a highly similar, but not identical, event to that which was falsely remembered actually did occur. This makes rejection of the false event (critical lure) at retrieval a demanding cognitive task. Given the prevalence of false memories in aging, it is important to identify means for mitigating them. The current study aims to examine the neural correlates underlying a retrieval-based monitoring strategy (RBMS; training older adults to monitor details of past events during memory retrieval in order to differentiate between true events (targets) and related but new events (critical lures) to reduce false memories in aging.

Age-related increases in false memories are most pronounced when targets share common features (e.g., perceptual elements, semantic labels) with lures (Balota et al., 1999; Kensinger & Schacter, 1999; Koutstaal & Schacter, 1997; Norman & Schacter, 1997; Schacter et al., 1997; Tun et al., 1998). Despite increased in false memory rates in aging, hit rates to targets in the same studies are typically equitable across age groups. This resulting behavioral pattern suggests that older adults may use general features of category membership to make their memory decisions, while not relying on encoded details to differentiate targets from related lures. Such target-lure differentiation requires not only memory for encoding-related details of target events, but also the ability to monitor for such details at the time of retrieval (Johnson & Raye, 1981; Lindsay & Johnson, 2000; Lyle & Johnson, 2007; Mitchell et al., 2000). While studies have shown that impaired retrieval monitoring contributes to age-related increases in false memories (Dodson & Schacter, 2002; Schacter et al., 1998), it is also important to note that research finds that older adults do, in fact, typically encode the details that are needed to support a distinction between targets and lures, yet fail to use them effectively during retrieval (Bowman & Dennis, 2015; Bulevich & Thomas, 2012; Cohn et al., 2008; Koutstaal, 2003; Mitchell et al., 2013; Multhaup, 1995; Park et al., 1984; Pezdek, 1987; Rahhal et al., 2002). For example, whereas older adults have shown greater false recognition of related lures on a standard old/new recognition task, they have shown equal performance to that of younger adults when utilizing detail-based memory in repetition priming and meaning-based recognition. This suggests older adults do not utilize details that are successfully encoded as effectively as younger adults (Koutstaal, 2003). Further, when older adults are provided specific instructions at retrieval to search for relevant perceptual and contextual cues when making their memory decisions, they are able to reduce their false memories, thereby improving overall memory discrimination (Bulevich & Thomas, 2012; Henkel, 2008; Koutstaal et al., 1999; Thomas & Bulevich, 2006). The fact that the instructions in the foregoing studies came after encoding, but prior to retrieval, also supports the notion that older adults encode, but fail to use, encoded details effectively during retrieval.

Strategy-based cognitive training has shown to be successful in enhancing true memories in older adults (Ball et al., 2002; Belleville et al., 2006, 2011; da Silva & Sunderland, 2010; Jennings & Jacoby, 2003; Kirchhoff et al., 2012, 2012b; Rebok et al., 2014; Willis, 1990). For example, following five days of adaptive training aimed at improving recollection, Jennings and Jacoby (2003) showed that older adults improved their ability to detect targets. Another study by Belleville and colleagues (2006) showed that following cognitive training via theoretical instruction and application to everyday life, both healthy older adults and older adults with mild cognitive impairment showed significant improvements on episodic memory tasks, including delayed list recall and face-name associations. Collectively, these findings suggest that specific strategy-based cognitive training can modify response criterion in older adults during memory retrieval.

In addition to the ability to modify behavior, neuroimaging studies have also shown that providing older adults with a memory strategy can improve performance through the modulation of neural processing during both encoding (Berry et al., 2010; Kirchhoff, Anderson, Barch, et al., 2012; Nyberg et al., 2003) and retrieval (Belleville et al., 2011; Hampstead et al., 2012; Kirchhoff et al., 2012b). For example, Kirchhoff and colleagues (2012b) found that semantic strategy training not only led to increased recognition memory performance in older adults, but was associated with training-related neural increases in bilateral hippocampus, bilateral middle and inferior frontal gyri, and right superior temporal cortex during retrieval. Further, activity within the medial superior frontal gyrus and left middle and inferior frontal gyri was also associated with self-initiated semantic strategy use during encoding (Kirchhoff, Anderson, Smith, et al., 2012), suggesting that brain activity changes were due to older adults’ increased use of semantic strategies that encourage the use of contextual information that enables recollection. Similar neural increases within the middle frontal gyrus and left inferior parietal lobe have also been found following mnemonic training (including techniques like semantic organization, semantic elaboration, and mental imagery) in aging (Belleville et al., 2011; Hampstead et al., 2012). Taken together, the results suggest that older adults are capable of utilizing training to engage effective memory strategies and that neural recruitment can be modulated to benefit memory performance in advanced aging. Yet, to date, such strategies have not been applied to the reduction of false memories.

Independent of memory, monitoring and cognitive control-based cognitive training has also been found to modulate prefrontal cortex functioning in aging (Basak et al., 2008; Braver et al., 2009; Braver & Barch, 2002). For example, Braver and colleagues (2009) enhanced proactive control in older adults by providing focused strategy-based cognitive training. While older adults showed inefficient increases in prefrontal cortex activity relative to younger adults prior to training, following training, they showed a similar pattern of neural recruitment to that of younger adults in lateral dorsolateral prefrontal cortex and the inferior frontal junction. In another example, Olesen and colleagues (2004) found increased activation in the middle frontal gyrus and superior and inferior parietal cortices following working memory training. These results suggest that strategy-based cognitive training in older adults can directly influence cognitive control processes via changes in prefrontal function. Overall, prior strategy training results suggest that older adults are capable of utilizing cognitive training to enhance memory performance and that this is accompanied by modulated neural recruitment that benefits memory performance in advanced aging.

Multivariate analyses may provide more detailed information about unregulated neural recruitment, showing that brain regions may exhibit changes in the discriminability of memory-related neural patterns, reflecting changes in their ability to behaviorally differentiate between targets and related lures following memory training. For example, neural patterns associated with perceptual categories become less discriminable within the ventral visual stream in the context of increasing age (Carp et al., 2011; Trelle et al., 2020). This dedifferentiation of neural patterns is related to measures of fluid processing abilities and memory performance within older adults (Koen et al., 2019; Koen & Rugg, 2019; Park et al., 2010; Sommer et al., 2019). Multivariate neuroimaging approaches, specifically multivariate classification, have been useful in assessing such differences in neural patterns associated with memory performance and how these patterns are altered by age. In an early example, Quamme and colleagues (2010), used multivariate classification analyses to demonstrate that neural patterns associated with recollection and familiarity within middle temporal gyrus correlated with correct rejection rates. Additionally, work from our group (Bowman et al., 2019) found neural patterns associated with target and lure items of varying relatedness were discriminable in portions of the ventral visual stream during memory retrieval, and that older adults exhibited an age-related reduction in neural discriminability in select portions of the ventral visual cortex. Interestingly, while younger adults exhibited consistent positive relationships between neural discriminability and memory discriminability (d’), older adults depicted a negative relationship in fusiform gyrus. Classification searchlight results suggested that regions outside the ventral visual stream, such as medial temporal, parietal, and frontal regions, may also maintain discriminable neural patterns associated with mnemonic information that are impacted by age. Collectively, prior reports suggest that subtle differences in information processing related to previously seen and unseen items are behaviorally relevant, and susceptible to age-related dedifferentiation. While behavioral work suggests that behavior discrimination is improved with cognitive training, no study has examined the impact of cognitive training on neural discriminability within the context of memory processing in aging. It may be that targeted cognitive interventions could influence both the magnitude of brain activity as well as the discriminability of neural information in cortical regions.

The current study aims to examine the cognitive and neural effects of retrieval-based monitoring strategy (RBMS) training. Based on previous findings, we hypothesize that RBMS training will lead to reduced false memories to related lures as well as modulation of the frontoparietal monitoring network at retrieval. Specifically, we posit that RBMS training will be associated with enhanced activity within the frontoparietal monitoring network. We further hypothesize that RBMS training will result in increased neural discriminability within the frontoparietal retrieval network, and that such increases will be associated with decreases in false memories.

## 2.0 Methods

### 2.1 Participants

Fifty native English-speaking older adults were recruited from Centre County, Pennsylvania. Participants received fMRI screening over the phone, including screening for neurological disorders and psychiatric illness, alcoholism, drug abuse, and learning disabilities. Once fMRI eligibility was determined, participants visited the lab and completed a battery of cognitive assessments. Subsequently, participants were pseudo-randomly assigned to active non-adaptive control or training groups balanced for age and gender. Participants completed written informed consent, which was approved by the Pennsylvania State University IRB committee and were compensated for their participation.

Of the 50 older adults who participated in the study, three dropped out due to illness or other personal reasons. Data from four participants were excluded due to technical difficulties/errors. Two participants were unable to provide scanner eligibility documentation. One participant had an incidental MRI finding and could not continue. Data from two participants were removed due to noncompliance with task instructions. As a result, our final analyses included complete data from 38 participants (age range=60-85 years old; mean age= 67.29; 15 males). Nineteen participants were in each group (see Table 1 for full cognitive assessment and demographic information for each group).

**Table 1.**
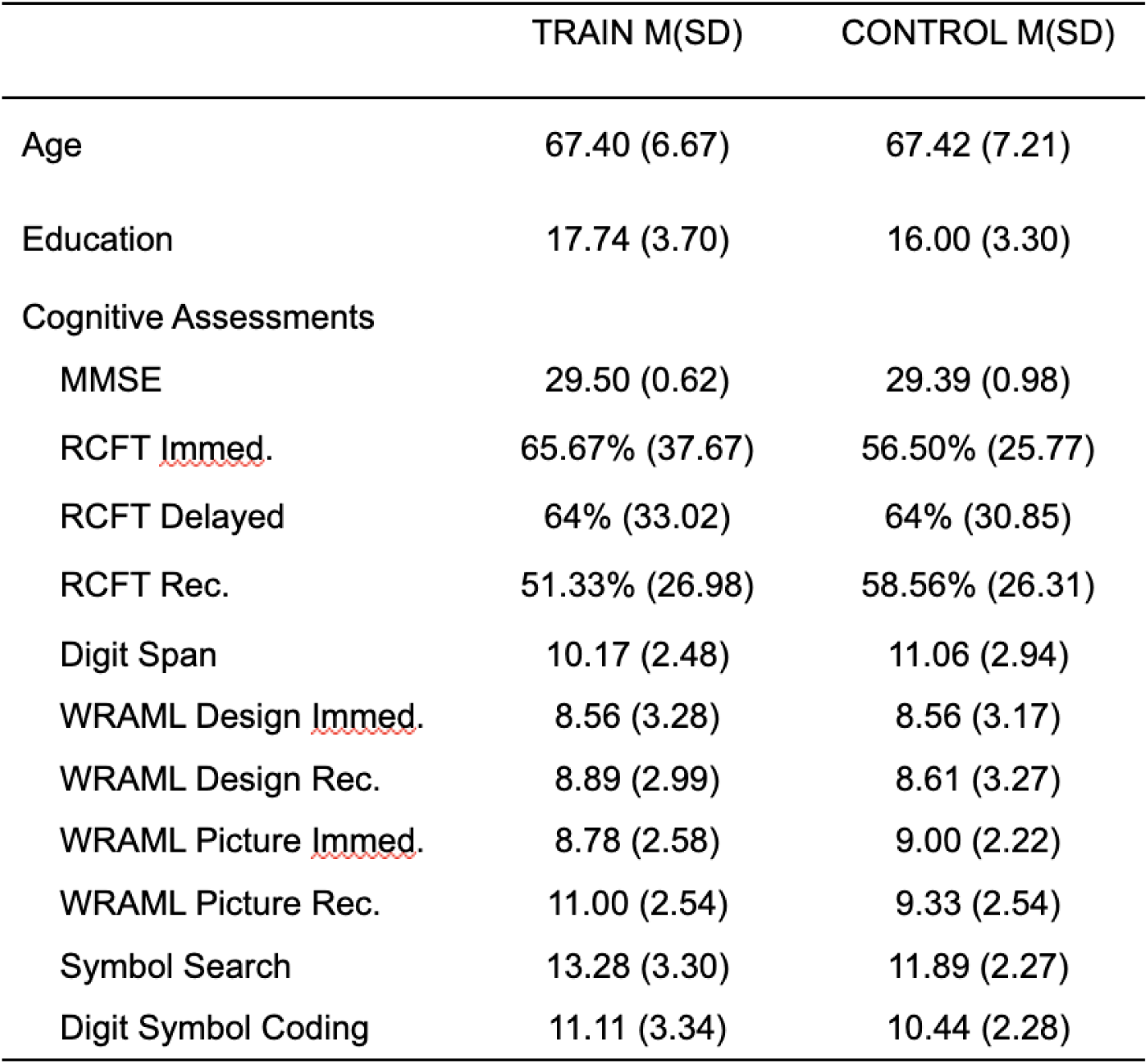
Demographical information of participants. M = Mean; SD = Standard Deviation; MMSE = Mini-Mental State Exam; RCFT = Rey Complex Figure Test; Immed. = Immediate; Rec. = Recognition. WRAML = Wide Range Assessment of Memory and Learning. All cognitive assessments were t-tested with Benjamini-Hochberg corrections, we observed no significant differences between groups.

### 2.2 Stimuli

Stimuli consisted of 804 color pictures of common objects gathered from internet searches, including those used in previous lab studies (Bowman & Dennis, 2015; Dennis et al., 2012). All backgrounds were removed, and pictures were cropped and resized to an approximate size of 480 × 480 pixels. Images were presented focally and equated for resolution. Images were displayed by COGENT in MATLAB (Mathworks). No categories were repeated during pre-training and post-training sessions.

### 2.3 Procedure

#### 2.3.1 Pre-Training and Post-Training fMRI Sessions

Procedures for “Pre-Training” and “Post-Training” days were identical (See Figure 2 for an overview of the study design). Older adults first performed an intentional encoding task outside of the MRI scanner where participants viewed a total of 120 images (40 categories, 3 exemplars per category) across two presentation blocks, each lasting approximately four minutes. Individual images were presented for 3 seconds on a black background followed by a 1500 ms interstimulus interval where a fixation cross was presented. Participants were asked to make a size judgement (Is the object smaller or bigger than a shoebox in real life?) for each image and to record their responses with a keypress. The images were pseudorandomly ordered to ensure that the three images from any given category did not appear consecutively. Following encoding there was a 20-minute retention interval during which instructions for the retrieval task were provided, participants entered the scanner, and structural images were acquired.

**Figure 2.**
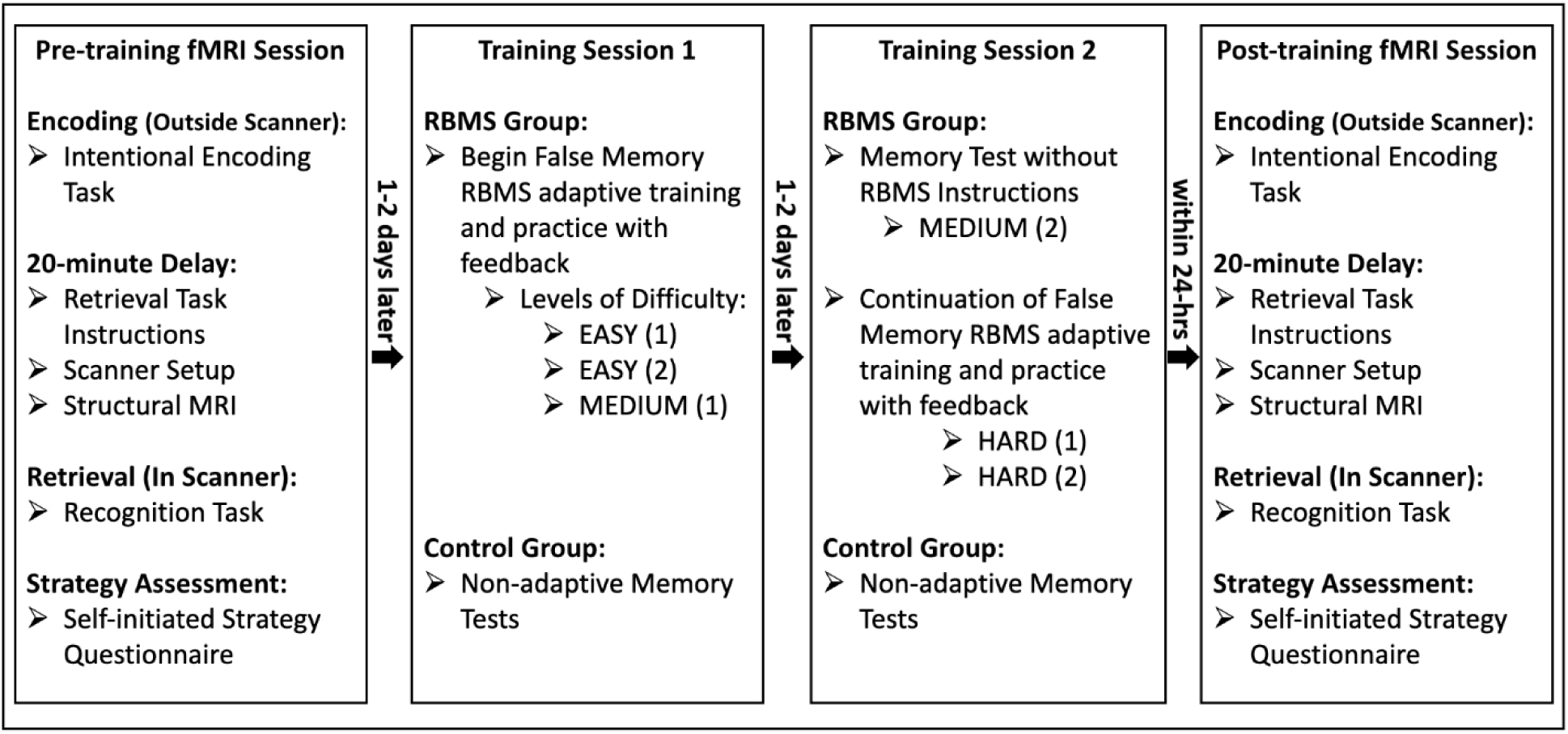
Study overview. During the pre- and post-training fMRI sessions, participants performed an intentional encoding task outside of the scanner, followed by a 20-minute interval delay. Subsequently, participants completed a recognition task during fMRI scanning. During training session 1, Participants in the control group completed non-adaptive memory tests, while participants in the training group received false memory warnings, along with adaptive Retrieval-Based Monitoring Strategy (RBMS) training. During training session 2, participants in the control group continued to complete non-adaptive memory tests, while participants in the training group continued to receive adaptive RBMS training according to their accuracy level. Finally, all participants completed an intentional encoding task followed by an fMRI recognition task.

During the memory retrieval task inside the MRI scanner, participants viewed 268 images (120 targets, 120 related lures, and 28 unrelated lures) across 4 runs, each lasting approximately 6 minutes. All images were presented in the center of the screen with a black background. Recognition response options with confidence ratings (“Old-High”, “Old-Low”, “New-Low”, “New-High) were displayed below each image. Behavioral responses were recorded using a 4-button response box. Participants were instructed that while some images would seem similar to those which were presented during the study phase, they should only respond ‘old’ if the exact image had been previously presented. Images were presented for 3 seconds, during which time participants made their memory responses to the given image. Image presentation was separated by a varied interstimulus interval with a fixation cross. The images were pseudorandomly presented to ensure that no more than three images from any one trial type (target, related lure, unrelated lure) appeared in a row. Including set up, structural scans, and the retrieval task, the total duration of scanner time for each participant was approximately 45 minutes.

Immediately after completing the memory retrieval task in the scanner, participants completed a paper and pencil strategy-use questionnaire outside of the scanner. The questionnaire asked about the strategies, if any, the participant used to try to remember items during the retrieval memory test. Questions included two questions: 1) During the **MEMORY TEST** task, did you try to remember the objects that were presented to you? And 2) If so, what strategy or strategies did you use to try to remember the objects?

During the post-training scan session, no mention of the training task was provided. The experimenter running the scanner during this session was blind to the participant’s group. Participants were debriefed at the end of the session (See Figure 1 for examples of stimuli).

**Figure 1.**
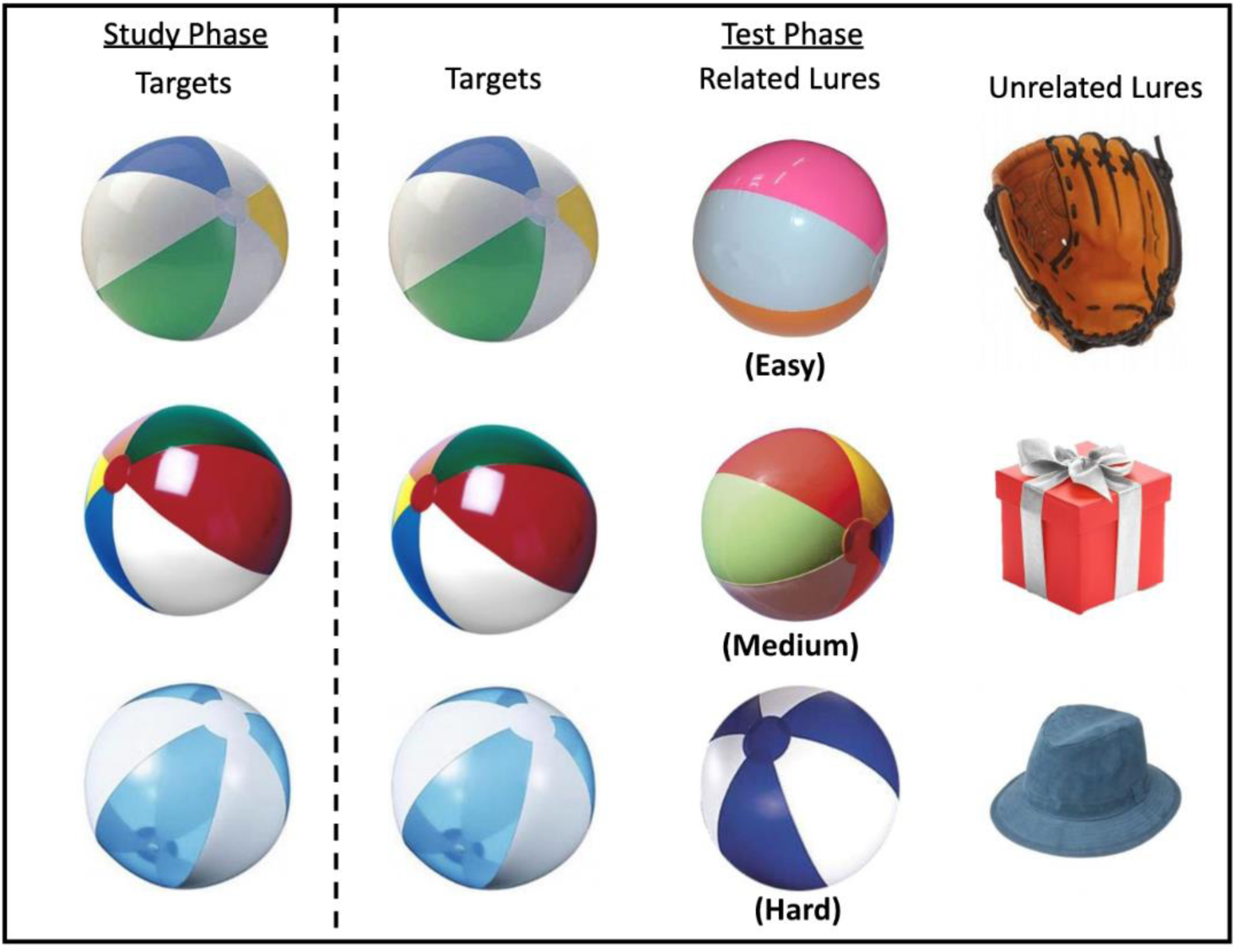
Example stimuli. Example of the study design that participants viewed during the fMRI and training sessions. Target items were shown at encoding followed by retrieval. At retrieval targets were shown, as well as related and unrelated lures. Examples of easy, medium, and hard related lures are shown.

#### 2.3.2 Training Sessions

Participants were randomized to either an active control or a retrieval-based monitoring strategy (RBMS) training condition by an independent researcher. The participant groups underwent either non-adaptive practice or RBMS training on two separate days after the pre-training neuroimaging session. Training was provided individually to each participant by researchers blinded about how the participant performed at baseline, before training. Participants in the non-adaptive practice group were given no instruction or emphasis on monitoring or paying attention to perceptual details.

During the first training session, participants in both groups first completed a DRM (Deese Roediger McDermott) task (Deese, 1959) in which they studied lists of semantically related words (e.g., nurse, hospital, etc.). During encoding, participants were instructed to listen to a series of words. After encoding, they completed a recognition memory task of the words where they are asked whether they remember the previously presented words, among related, but not presented words (e.g., doctor).

Participants in the RBMS training group were then given a brief overview of the theoretical basis of false memories. Participants were informed of the definition of a false memory and were provided with an explanation of how they may occur during a memory task. Participants were then given training aimed at helping them evaluate and monitor, during retrieval, the specific details of items from encoding so that they could distinguish whether an item was the same as one which was seen at encoding, or merely similar to it. For example, they were told that focusing on details such as “a small, black, fluffy dog with brown spots” during encoding would help them make a correct memory decision during retrieval when presented with either the target dog or a similar small dog. Participants in the RBMS training group were instructed on how to search their memory and accomplish this strategy. It was explained that solely retrieving at a superficial, gist level (e.g., small dog) had the potential to lead to false memories and erroneous endorsement of a critical lure, such as a different, but similar dog. The use of these instructions and training were implemented first in simple perceptual discrimination tasks, and then for memory tasks. At the end of each task, participants received accuracy feedback (i.e., how well they performed) and were asked to explain to the experimenter how monitoring item-specific details helped their performance.

RBMS training was adaptive and gradually increased in difficulty as participants’ performance improved. Specifically, participants in the RBMS group completed memory tests at two EASY levels, EASY (1) & EASY (2), followed by MEDIUM (1), MEDIUM (2), HARD (1), and HARD (2) difficulty levels (different stimuli were used in each test). Participants repeated a level if their hit rate and/or false alarm rate was poor (hit rate below 60% or false alarm rate above 60%). Figure 3 presents the number of stimuli during encoding and retrieval, the number of categories, the number of stimuli per category for targets and lures, the number of unrelated lures, the difficulty of stimulus discriminability, and the retention interval for each RBMS and non-adaptive training trial. The total duration of the first training session was approximately 1.5 hours. The first training session always concluded with either a MEDIUM (1) or MEDIUM (2) level.

**Figure 3.**
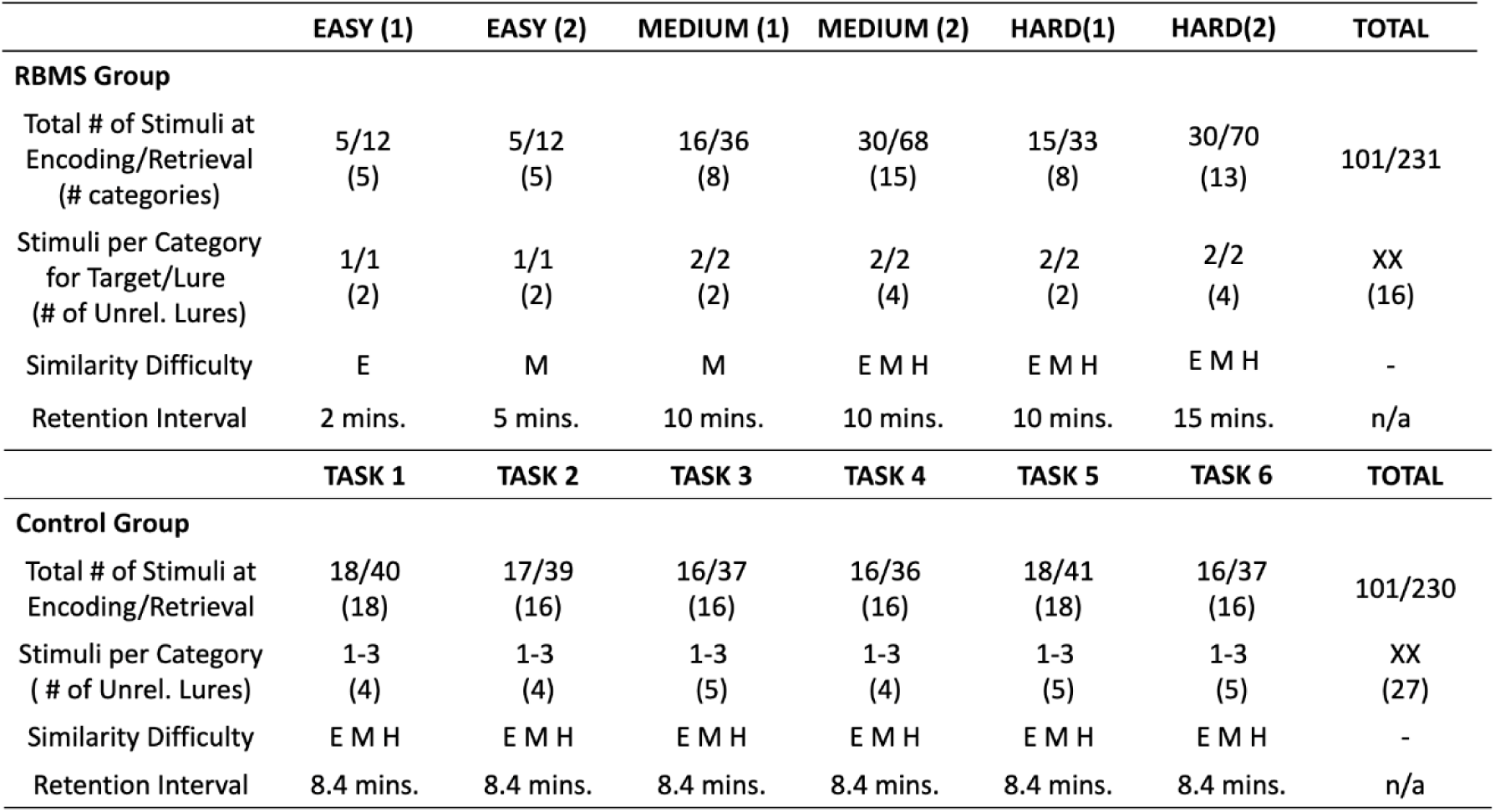
Retrieval-based Monitoring Strategy Training Design. Training was adaptive with feedback, gradually increasing in difficulty (i.e., EASY (1), EASY (2), MEDIUM (1), MEDIUM (2), HARD(1), HARD (2)), as participant’s performance increases, by adding multiple exemplars within a category at encoding and retrieval, increasing the retention interval (time between encoding and retrieval), as well as varying item distinctiveness or similarity: E = Easy, M = Medium, H = Hard. Unlike the RBMS group, the control group received 6 memory tasks, including all stimuli from the RBMS training phases. These tasks were not adaptable in nature, instead, in all 5 tasks, participants viewed 1 to 3 stimuli per category, stimuli included images from all 3 levels of the item distinctiveness, and retention interval were 8.4 seconds, which is the average retention interval to be completed by the RBMS training group. Unrel. = Unrelated.

During the second training session, participants in the RBMS training group completed more memory training. A goal of the second training day was also to assess how well each participant was able to spontaneously use the retrieval-based monitoring strategy. Therefore, for the first memory test given on training day 2, the participant was not instructed use the retrieval-based monitoring strategy. Instead, they completed a brief memory test trial and were given memory task instructions identical to those used during the pre-training fMRI scanning session. Following this first memory test, the participant was reminded of the importance of using the retrieval-based monitoring strategy and was re-instructed on how to execute it using examples from their own training session 1 training task performance. Training then resumed at the next MEDIUIM difficultly level following where the participant left at the end of their first training session. Training subsequently continued through the HARD training tests. The second training session ended with another DRM task trial. The total duration of the second training session was approximately 1 hour and 30 minutes.

Across both training days, the active control group completed multiple memory tests that used stimuli identical to those used for the RBMS training group (Table 3). The active control group spent an equal amount of time in the lab completing memory tests as the RBMS training group. However, no information about false memories or retrieval monitoring was provided to active control group participants. Instead, they were given memory task instructions identical to those used during the pre-training fMRI scanning session. Active control group participants were told that practice on memory tasks was known to lead to improved performance, so therefore they would be completing many practice trials/sessions.

**Table 3.**
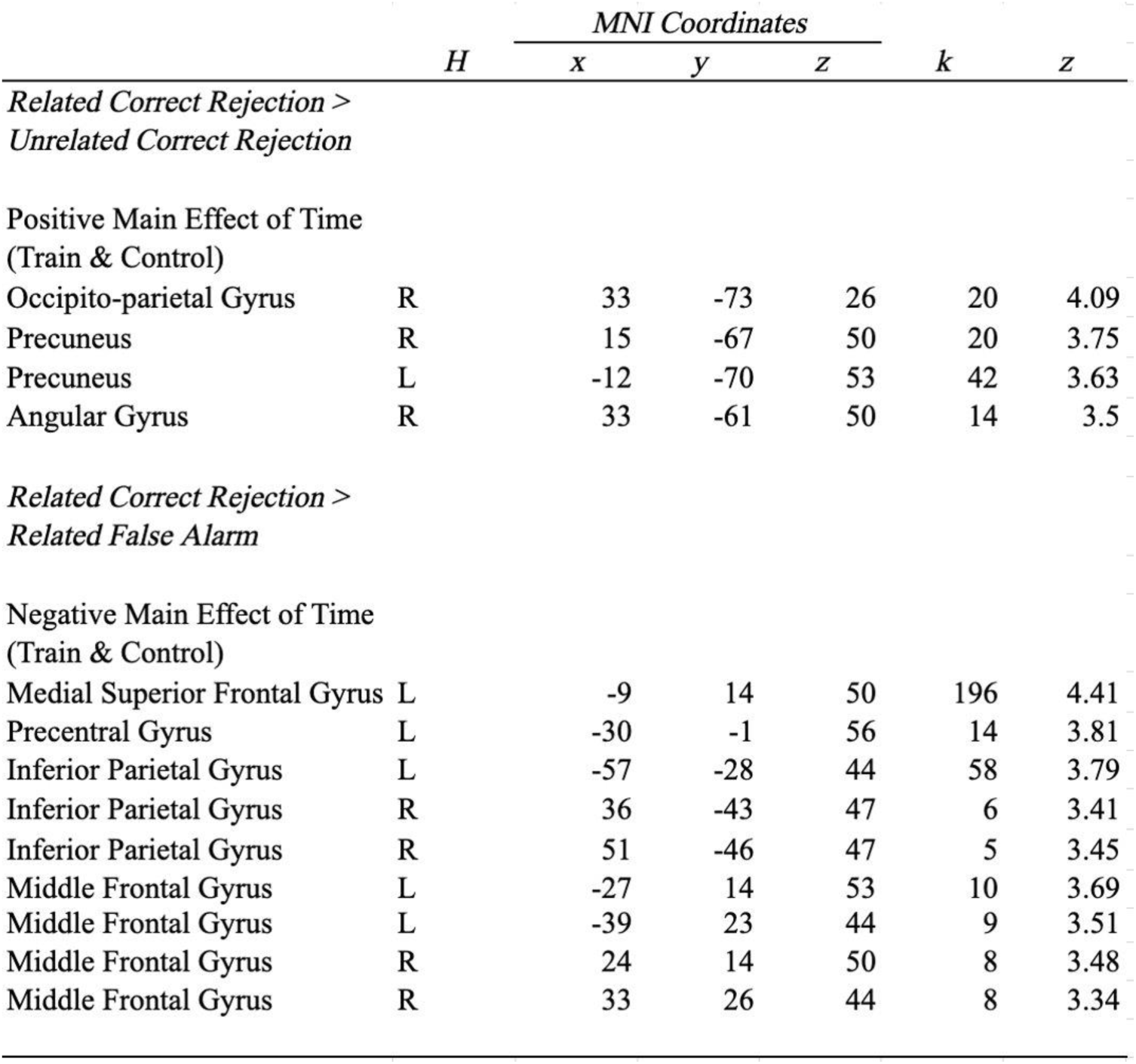
The table reports Pre-Post Effects on Univariate Activity for general monitoring (Related Correct Rejections > Unrelated Correct Rejections) and successful monitoring (Related Correct Rejections > Related False Alarms). X, y, z represents peak MNI coordinates, k indicates cluster extent, H represents hemisphere, L – left, R-right, z – statistic.

### 2.4 Image Acquisition

Structural and functional brain images were acquired using a Siemens 3T scanner equipped with a 12-channel head coil. A T1-weighted sagittal localizer was collected to locate the anterior (AC) and posterior (PC) commissures. A high-resolution anatomical image was then acquired with a 1650ms TR, 2.03ms TE, 256mm field of view (FOV), and 2562 matrix with 160 1mm thick axial slices resulting in 1mm isotropic voxels. Echo-planar functional images were acquired using a descending acquisition scan with a 2500ms TR, 25ms TE, 240mm FOV, and 802 matrix with 42 3mm thick axial slices resulting in 3mm isotropic voxels.

### 2.5 Behavioral Analyses

We assessed potential demographic characteristics at baseline assessment via two-sample two-sided t-tests. Multiple linear regression was used to examine group differences in memory performance (Hits and False Alarms) at time point 2 (T2; Post-Training fMRI Scanning Session). Memory performance at time point 1 (T1; Pre-Training fMRI Scanning Session) and training group (RBMS, Control) were predictors.

### 2.6 fMRI Analysis

#### 2.6.1 Imaging data preprocessing

Pre-processing of all functional images was carried out in SPM12 (Wellcome Institute of Cognitive Neurology, London, UK. www.fil.ion.ucl.ac.uk) using MATLAB (Mathworks Inc., Natick, MA, USA). The functional time series were first corrected for differences in slice timing acquisition. EPI images were then realigned to the first image of the functional run using a 6-parameter rigid body affine transformation and then spatially normalized to the standard MNI (Montreal Neurological Institute) EPI template implemented in SPM12. To do this, the raw T1 MPRAGE images were co-registered to the mean realigned functional image, and then this co-registered T1 MPRAGE image was segmented and registered to the MNI template. Lastly, the parameters from this registration process were applied to the slice time corrected and realigned functional images to normalize them to the MNI template. As a final preprocessing step, all of the normalized functional images were smoothed using a 6mm full-width-half-maximum Gaussian smoothing kernel. Normalized unsmoothed data were used for multivariate classification analyses.

#### 2.6.2 Univariate analysis

At the first level, trial-related activity was modeled using the general linear model (GLM) with a stick function corresponding to trial onset convolved with a canonical hemodynamic response function. A second-level random effects GLM was created and one sample t-tests were conducted to investigate contrasts of interest. The data was sorted into the follow regressors: 1) All Hits, which were defined as both ‘Definitely Old’ and ‘Probably Old’ responses to related targets; 2) All Misses, which were defined as ‘Definitely New and ‘Probably New’ responses to related targets; 3) Related False Alarms (RFA), which were defined as both ‘Definitely Old’ and ‘Probably Old’ responses to related lures; and 4) Related Correct Rejections (RCR), which were defined as ‘Definitely New and ‘Probably New’ responses to related lures; 5) Unrelated Correct Rejections (UCR), which were defined as ‘Definitely New and ‘Probably New’ responses to unrelated lures; and 6) Unrelated False Alarms (UFA), which were defined as both ‘Definitely Old’ and ‘Probably Old’ responses to unrelated lures. All no response trials were coded with their own regressors and treated as regressors of no interest, as were movement parameters.

To examine brain regions that supported general monitoring, we contrasted related correct rejections with unrelated correct rejections. To examine successful monitoring, we contrasted related correct rejections with related false alarms. Both contrasts were conducted on pre-training fMRI data (T1) in order to identify regions of interest for conducting subsequent group analyses of training-related changes. We utilized a cluster-forming threshold of *p* < .001 with a voxel extent of 36 to determine significant clusters at the group level as determined by Monte Carlo simulations implemented in AFNI (Cox & Hyde, 1997). In cases where widespread univariate activity was observed precluding the identification of discrete significant clusters (See Results, Successful Monitoring), we masked the contrast map with theoretically relevant regions via the WFU PickAtlas (superior medial frontal gyrus, middle frontal gyrus, inferior parietal gyrus, superior and middle temporal gyrus, and fusiform gyrus) (Dennis et al., 2007; Kurkela & Dennis, 2016; Yu et al., 2019).

To examine neural changes associated with cognitive training, we submitted the first level general monitoring and successful monitoring contrast maps at T1 and T2 to the Sandwich Estimator (SwE version 2.2.2) toolbox implemented in SPM (Guillaume et al., 2014); http://www.nisox.org/Software/SwE/), masking for regions that survived cluster correction at the group level (N = 38) at T1 (see *Results*, Figure 4). The SwE toolbox applied non-iterative marginal models to the dataset while also accounting for correlations due to repeated measurements and error variation across individual participants. The toolbox is well adapted to handle datasets that are small or potentially unbalanced. Specifically, we examined effects of time (pre-training to post-training) in the full sample as well as in the RBMS and control groups separately. Noted above, we limited our analysis to regions that displayed significant differential BOLD amplitude at the second level in the full sample during the pre-training fMRI session. This allowed us to examine changes in neural activity associated with general monitoring and accurate monitoring in processing-relevant regions. We used a cluster-forming threshold of *p* < .001 and k = 4 as determined by Monte Carlo simulations implemented in AFNI (Cox & Hyde, 1997) to identify significant pre-/post-findings. Specifically, we first examined any potential group differences in univariate activity at post-test while controlling for univariate activity at baseline. We then examined potential time-related changes in univariate activity. We next conducted several exploratory regression analyses. Specifically, we used change in univariate contrast estimates between T2 and T1 as the outcome variable, and included group, contrast estimate at T1, and their interaction as predictor variables. We also examined if any changes in neural activity contributed to changes in correct rejection rates in older adults by conducting multiple linear regressions in which change in correct rejection rates (T2 minus T1) was the outcome variable and group, change in contrast estimate, and their interaction were predictor variables.

**Figure 4.**
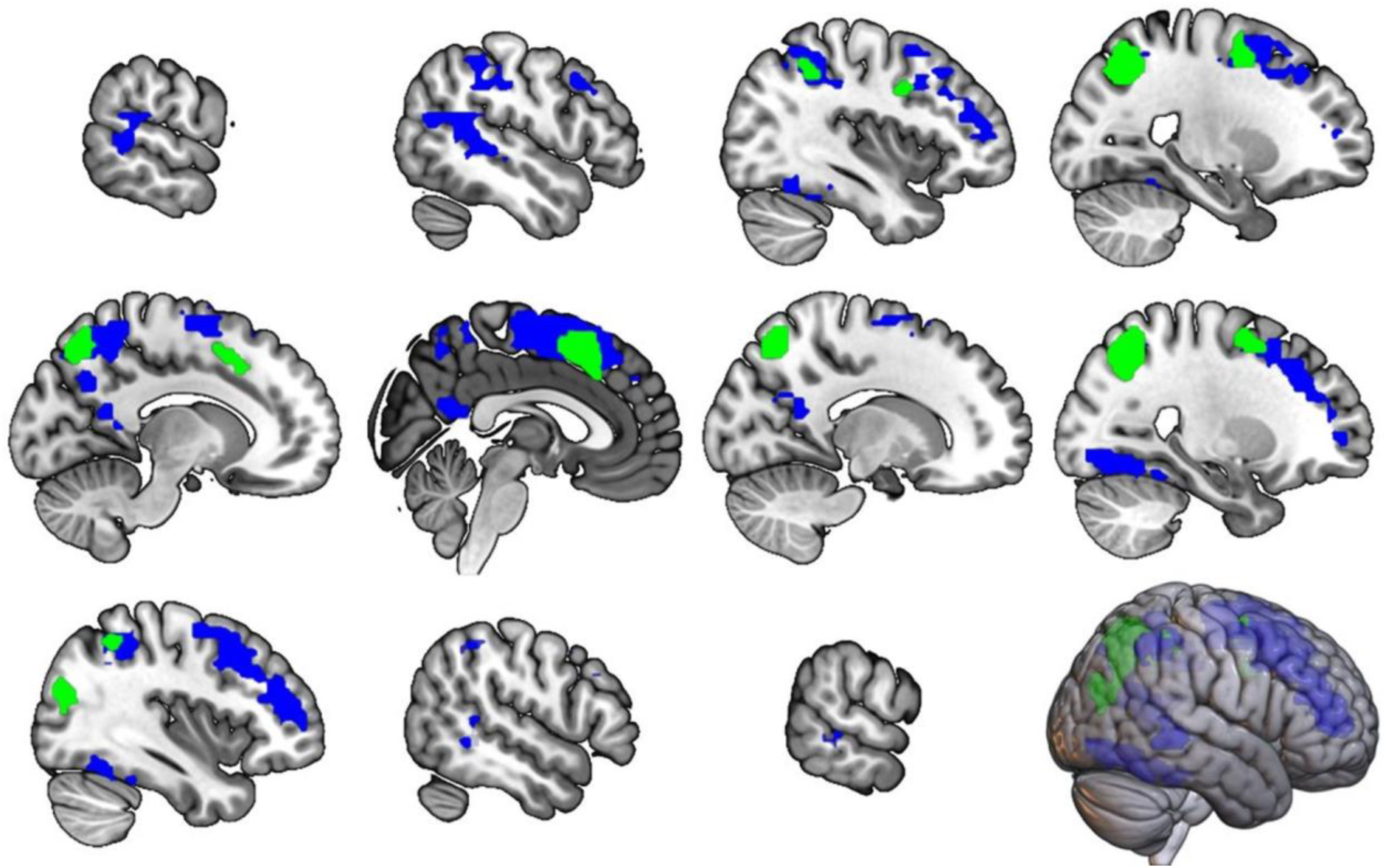
Baseline univariate activity. Regions that survived cluster correction at the group level (N=38) at T1 for general monitoring (Related CR vs Unrelated CR) in green, and successful monitoring (Related CR vs Related FA) in blue.

#### 2.6.3 Multivariate Classification Analysis

In addition to examining univariate changes in neural activity following training, we were also interested in examining training-related changes in neural discriminability and how that may relate to behavioral improvements in older adults. To do so, we first estimated a second GLM in SPM12 defining one regressor per trial during memory retrieval with normalized unsmoothed data (Op de Beeck, 2010). We included six additional nuisance regressors for head motion. We estimated whole-brain beta parameter maps for each trial and for each participant. We then concatenated each beta parameter map across each retrieval run and submitted this data to classification analyses using the CoSMoMVPA toolbox (Oosterhof et al., 2016). Specifically, we conducted classification analyses distinguishing between patterns of neural activity underlying target and related lure items (Bowman et al., 2019). Regions of interest included regions identified as significant in the monitoring contrasts at T1, with any overlapping clusters collapsed to create a single ROI (see Supplemental Figure 1). Critically, the ROIs were defined using contrasts examining lure-related memory responses, specifically correct rejections and false alarms, whereas the multivariate classification analysis utilized targets and lures, thereby negating potential circular analysis confounds. We utilized a support vector machine (SVM) classifier with a linear kernel and all voxels within each region of interest. We used a leave-one-out cross-validation approach in which the classifier was trained on three runs and tested on one run. We averaged across validation folds from all possible train/test permutations to estimate subject-level classification accuracy. We first conducted classification analyses in each region of interest, in each participant, at pre-test. To test if the SVM classifier was able to discriminate neural patterns associated with targets and lures above chance (50% accuracy), we used one-tailed one-sample t-tests in each region at T1 collapsing across the train and practice groups. We then repeated the above classification procedure for all participants and ROIs at T2, examining classifier performance in each group separately. To examine possible group differences in neural discriminability at T2 within regions that depicted significant above-chance classification accuracy while controlling for baseline discriminability, we conducted primary multiple linear regression analyses in which classification accuracy at T2 was the outcome variable, and group, classification accuracy at T1, and their interaction were predictor variables. We also conducted exploratory regression analyses in regions depicting above-chance classification. Specifically, change in accuracy between T2 and T1 was the outcome variable, and group, classification accuracy at T1, and their interaction were predictor variables. Finally, we wished to examine if any changes in neural discriminability contributed to changes in correct rejection rates in older adults. We therefore conducted multiple linear regressions in which change in correct rejection rates (T2 minus T1) was the outcome variable, and group, change in classification accuracy, and their interaction were predictor variablesany significance was confirmed with using a linear permutation model (lmPerm 2.1.0 package) of 10,000 permutations.

## 3.0 Results

### 3.1 Demographics and Cognitive Assessment

The sample included 19 control and 19 RBMS training participants. Demographic details of the 38 participants included in the analysis, separated by training group, are shown in Table 1. Participants were similar in age and years of education across training groups. Additionally, we observed no significant differences between training and control groups in any of the cognitive assessment tasks (all *p*’s > .05).

### 3.2 Effects of RBMS Training on Hit and False Alarm Rates

To investigate the effectiveness of RBMS training, multiple linear regression was conducted to determine whether training group (RBMS, Control) and/or memory performance at T1 predicted change in memory performance (Hits and False Alarms) between T1 and T2. Group (Beta = -0.076, *p* = 0.038) and false memory rates at T1 (Beta = -0.308, *p* = 0.021) significantly predicted decreased rates of false memories from T1 to T2. False memory rates for the RBMS group at T1 (M = 0.501, SD = 0.123) were comparable to the control group (M = 0.573, SD = 0.150), (*t(*36) = 1.603, *p* = .118). False memory rates for the RBMS group at T2 (M = 0.385, SD = 0.098) were significantly less than that of the control group (M = 0.511, SD = 0.173), (*t*(36) = 2.757, *p* = .009). With respect to hit rates, Group did not significantly predict increases in hit rates at T2 (Beta = 0.034, *p* = .242). Hit Rates at T1 significantly predicted Hit rates at T2 (Beta = 0.365, *p* = 0.006). Hit Rates for the RBMS group at T1 (M = 0.729, SD = 0.100) were comparable to the control group (M = 0.777, SD = 0.126), (*t*(36) = 1.323, *p* = .194). Hit Rates for the RBMS group at T2 (M = 0.738, SD = 0.105) were not significantly different compared to the control group (M = 0.802, SD = 0.115), (*t*(36) = 1.799, *p* = .080).

### 3.3 Univariate Activity

#### 3.3.1 General monitoring

At T1, general monitoring activity was observed, collapsed across all participants, in the precuneus, middle frontal gyrus, precentral gyrus, precentral gyrus, and superior parietal and medial frontal gyrus (Table 2, Figure 4).

**Table 2.**
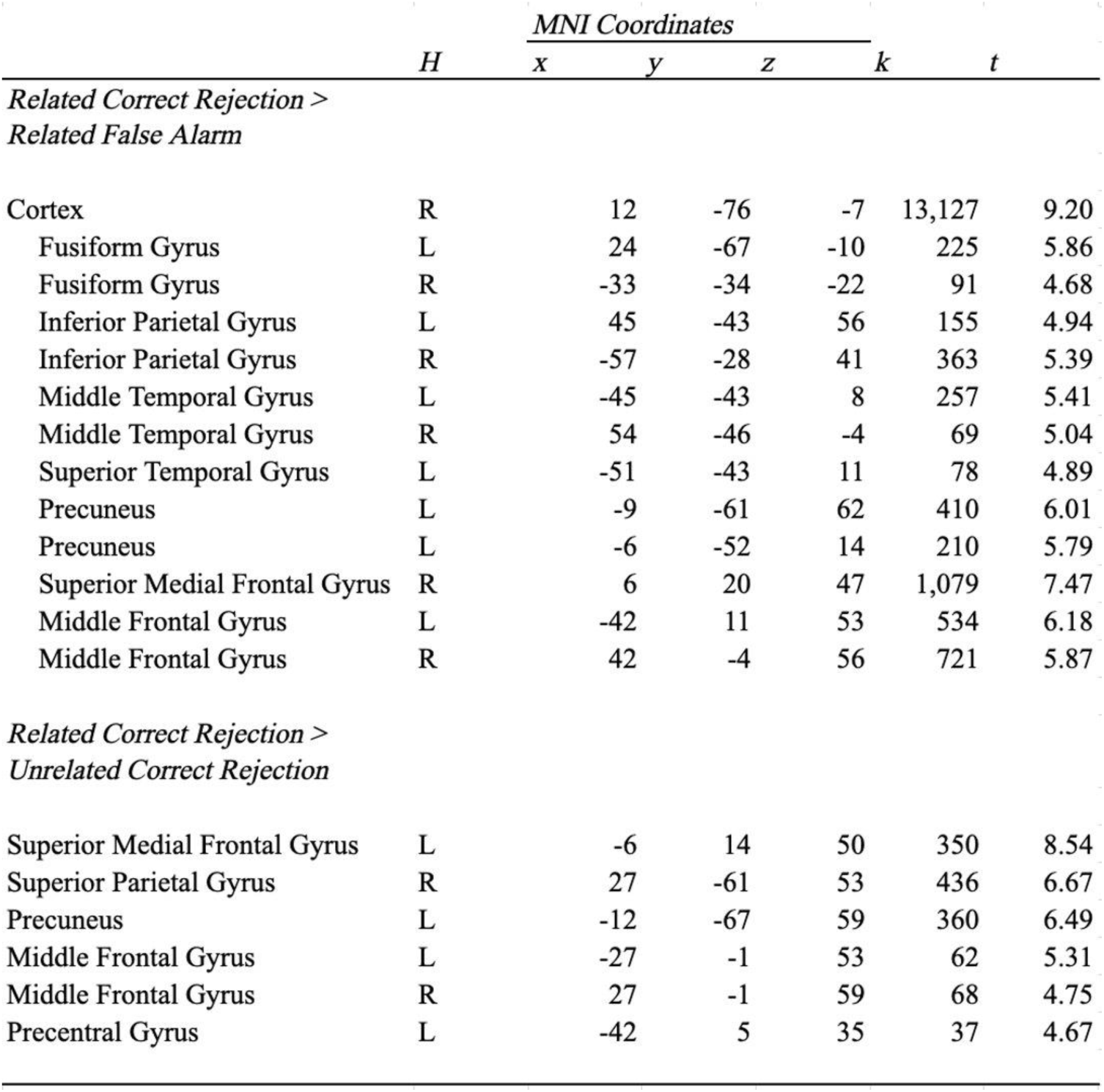
The table reports univariate activity for general monitoring (Related Correct Rejections > Unrelated Correct Rejections) and successful monitoring (Related Correct Rejections > Related False Alarms) in the full sample at baseline. X, y, z represents peak MNI coordinates, k indicates cluster extent, H represents hemisphere, L – left, R-right, t – statistic.

The results of the SWE analysis on the contrast of related correct rejections > unrelated correct rejections revealed a negative significant main effect of time reflecting a decrease in contrast estimates from T1 to T2 with regard to activation differences in several brain regions, including the precuneus, angular gyrus, and occipital-parietal gyrus (Table 3, Figure 5). There were no significant main effects or interactions associated with Group (See Supplemental Table 1 for results within each group separately). As an exploratory analysis, we examined if change in general monitoring contrast estimates could be predicted by group assignment or baseline performance with multiple linear regression. We observed no significant predictors of change in contrast estimates (all *p*’s > 0.05). We also examined if change in contrast estimates were predictive of change in correct rejection rates and observed no significant effects (all *p*’s > 0.05)

**Figure 5.**
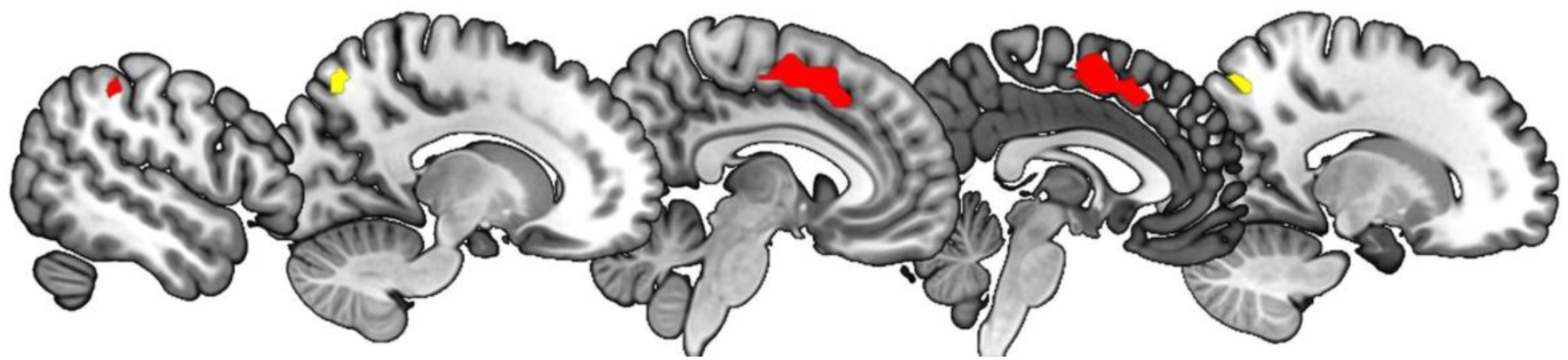
Pre-post changes in univariate activity. General monitoring seen in yellow: increase (Related Correct Rejections vs Unrelated Correct Rejections) and successful monitoring seen in red: decrease (Related Correct Rejections vs Related False Alarms). Slice numbers: -52, -14, -6, -2, 16

#### 3.3.2 Successful monitoring

At T1, successful monitoring activity was observed in widespread regions across the cortex. We therefore masked whole-brain results with theoretically relevant, structurally-defined regions to more precisely characterize the extent of activation, specifically the superior medial frontal gyrus, middle frontal gyrus, inferior parietal gyrus, superior and middle temporal gyrus, and fusiform gyrus. Successful monitoring activity was observed in the fusiform gyrus, inferior parietal gyrus, middle and superior temporal gyri, precuneus, middle frontal gyrus, and superior medial frontal gyrus (Table 2, Figure 4).

The results of the SWE univariate analysis on the contrast of correct rejections > related false alarms revealed increased neural activity from T1 to T2 in several brain regions, including the medial superior frontal gyrus, middle frontal gyrus, precentral gyrus, and inferior parietal gyrus

(Table 3, Figure 5). There were no significant main effects or interactions associated with Group (See Supplemental Table 1 for results within each group separately). As an exploratory analysis, we examined if change in successful monitoring contrast estimates could be predicted by group assignment or baseline performance with multiple linear regression. We observed no significant predictors of change in contrast estimates (all *p*’s > 0.05). We also examined if change in contrast estimates were predictive of change in correct rejection rates and observed no significant effects (all *p*’s > 0.05).

### 3.4 Multivariate Pattern Classification

We next examined whether brain activity patterns associated with targets and lures were discriminable in regions defined by the pre-training (T1) full-sample univariate analysis, and if neural discriminability in the regions was altered by cognitive training. The inferior parietal gyrus and the medial superior frontal gyrus displayed classification accuracies greater than chance (*t*(37) = 2.44, *p* = 0.010*; t*(37) = 2.06, *p* = 0.023, respectively) in the full sample pre-training. We next examined classification accuracy during the post-training fMRI session in the full sample. Again, both the inferior parietal gyrus (*t*(37) = 4.14, *p <* 0.001) and the medial superior frontal gyrus (*t*(37) = 2.64, *p* = 0.006), along with temporal cortex (*t*(37) = 3.29, *p* = 0.001) and the middle frontal gyrus (*t*(37) = 3.35, *p* = 0.001), displayed classification accuracies greater than chance (Figure 6).

**Figure 6.**
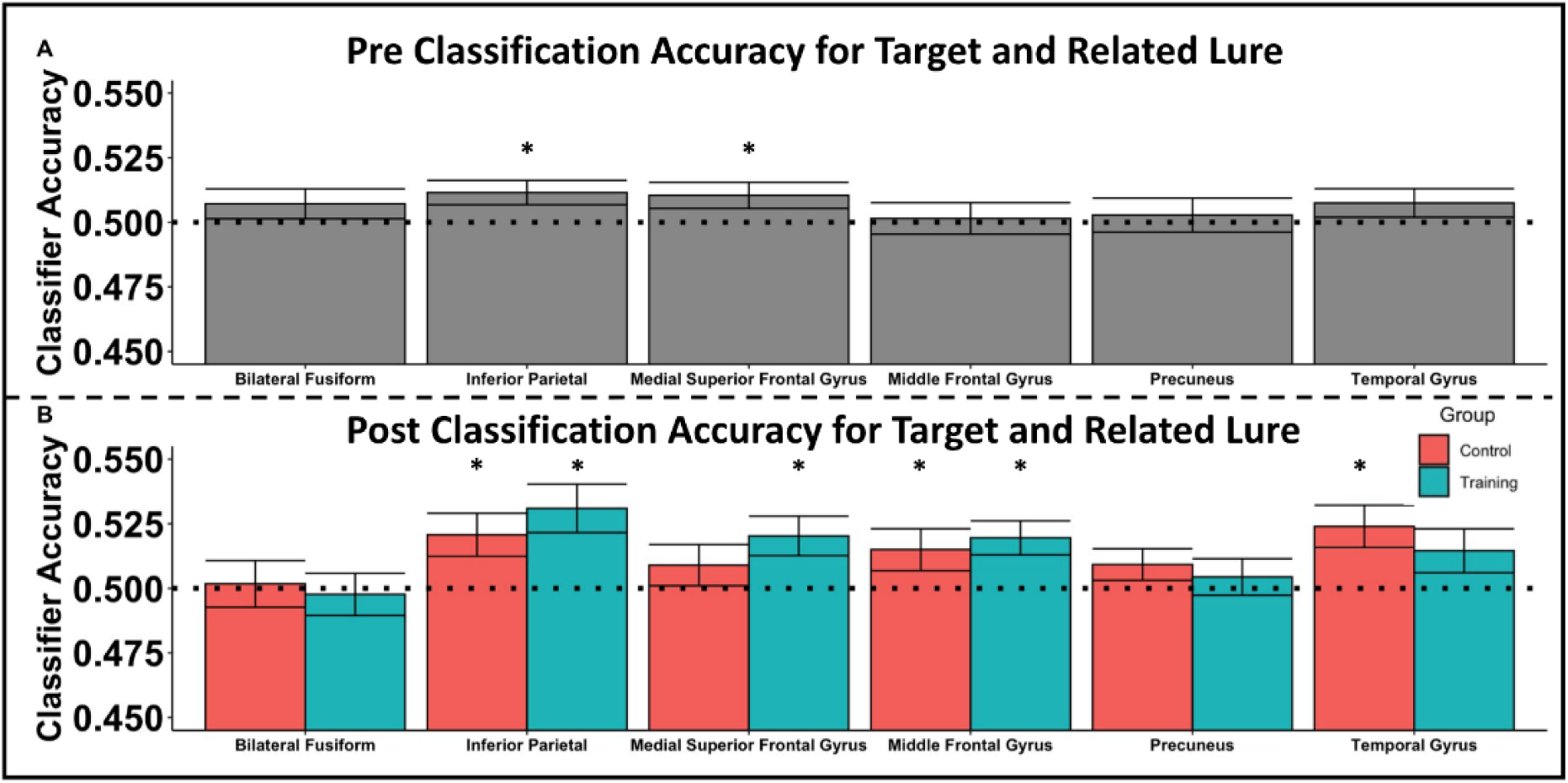
Multivoxel classification accuracy. A) Pre classification accuracy for targets and related lures of each of the regions of interest. * indicates regions that were significantly above chance (50%). B) Post classification accuracy for targets and related lures by group. * indicates the region and group that were significantly above chance (50%).

We also entered classification accuracy scores into multiple linear regressions to examine if there were any group changes in post-training classification accuracy when accounting for pre-training classification accuracy. Group assignment did not predict post-training classification accuracy within any ROI (all *p’s* > 0.05). We next ran multiple linear regression models with change-in-accuracy (post-training minus pre-training) as the outcome variable. We observed no significant group or group-by-pre-training interactions (all *p’s* > 0.05). However, we observed that pre-training (T1) classification accuracy significantly negatively predicted change in classification accuracy in several regions, including the inferior parietal gyrus (b(34) = -0.96, *p* = 0.005), middle frontal gyrus (b(34) = -1.15, *p < 0*.001), temporal cortex (b(34) = -0.59, *p* = 0.040), and precuneus (b(34) = -0.92, *p <* 0.001). In these regions, individuals with the lowest pre-training classification accuracy showed the greatest increase in classification accuracy over time (Figure 7). We next conducted exploratory multiple linear regression analyses that examined if changes in correct rejection rates over time were predicted by changes in classification accuracy. We observed no significant effects (all *p*’s > 0.05). We observed no changes in significance after conducting permutation testing.

**Figure 7.**
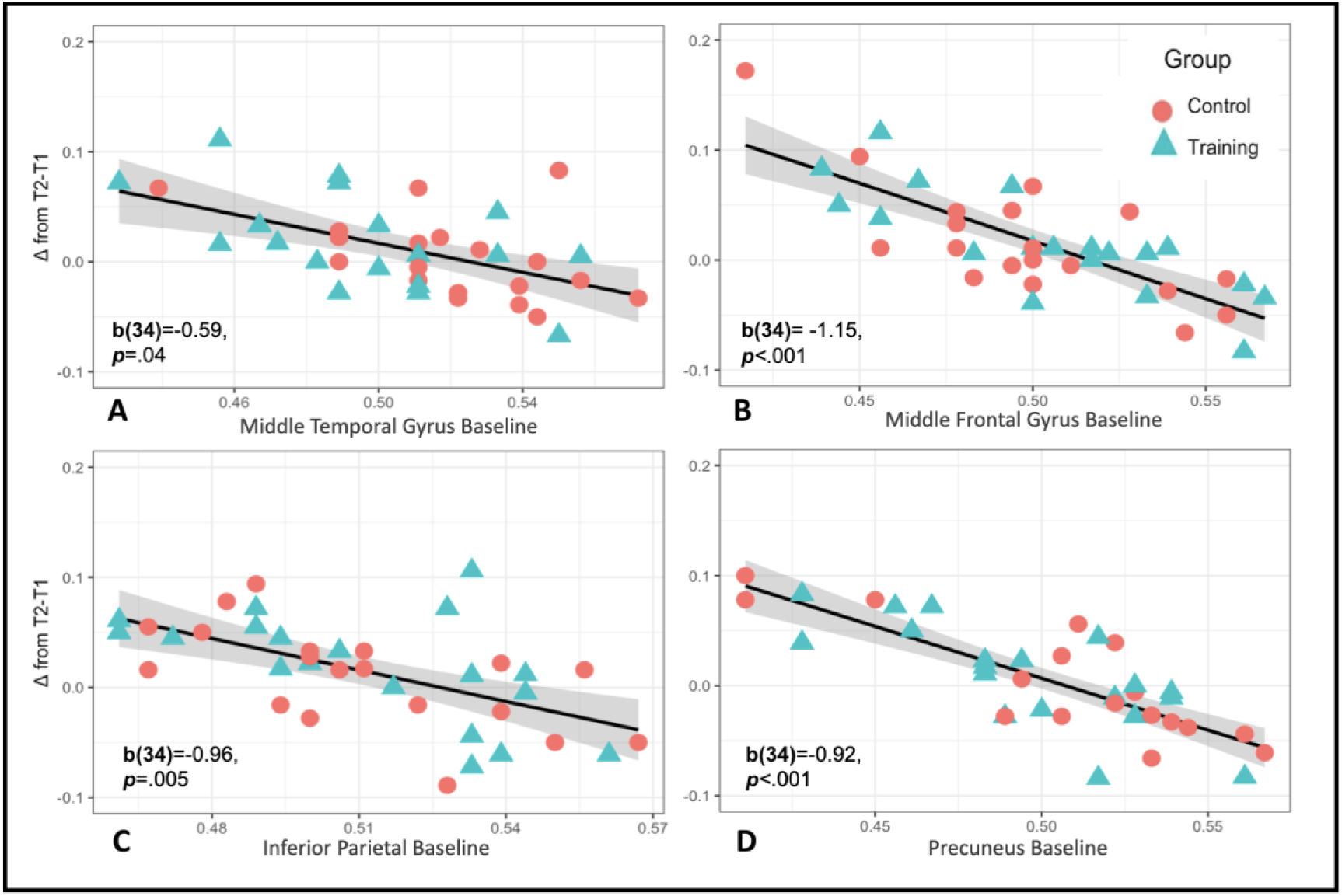
Change in classification accuracy. Scatterplots depicting change in classification accuracy (T2 minus T1) versus baseline classification accuracy in the A) middle temporal gyrus, B) middle frontal gyrus, C) inferior parietal gyrus, and D) precuneus. B= beta value, p= p-value.

## 4.0 Discussion

The goal of the current study was to investigate the cognitive and neural basis of a retrieval-based monitoring strategy (RBMS) training aimed at reducing false memories. Specifically, older adults were trained to use details of past events during memory retrieval to distinguish targets from related lures, with the goal of reducing false memories. As predicted, compared to participants in our control training group, participants in the RBMS training group showed a significant decrease in rates of false alarms following the intervention training. Additionally, we examined both univariate and multivariate approaches to understand the underlying neural processing pre- and post-training. Neuroimaging results revealed modulation of the BOLD signal within multiple frontoparietal regions associated with successful lure discrimination, as well as evidence of benefits to neural discriminability post-training within these regions. Together, these findings provide evidence supporting that training improved older adults’ ability to monitor distinguishing details of targets and lures during retrieval, with changes observed both behaviorally and neurally.

As mentioned earlier, older adults are prone to false recognition and misattribution errors. This is due, in part, to their reliance on general familiarity, as well as their decreased ability to retrieve specific recollections of the encoding event to either accept or reject a related lure presented during retrieval. Such failures in ‘recall to reject’ have been documented in both behavioral and neuroimaging work in aging (Cohn et al., 2008; Gallo et al., 2006; Yassa et al., 2001). Nevertheless, previous studies also suggest that older adults do encode sufficient details to distinguish between targets and lures, but they are unable to self-initiate a strategy to bring these details back online (Koutstaal, 2003; Koutstaal et al., 1999). Focusing on these points, the RBMS training encouraged participants to specifically use encoded details during retrieval to determine whether the presented items were previously encountered (i.e., old) or new. While the RBMS training significantly reduced rates of false alarms in our training group compared to controls, there was no change in rates of true memories between groups. This finding demonstrates that the RBMS training lends specifically to the discrimination and identification of new items presented during retrieval. Prior memory training and strategy intervention studies have seen improvements in older adults’ hit rates and behavioral discriminability (Ball et al., 2002; Belleville et al., 2006, 2011; da Silva & Sunderland, 2010; Jennings & Jacoby, 2003; Kirchhoff et al., 2012, 2012b; Rebok et al., 2014). However, this study stands among the first training studies to focus specifically on reducing false memories and to show reductions in errors of commission. As such, the training offers a unique contribution to methods aimed at improving age-related memory errors.

In addition to the observed decrease in false alarms, the RBMS group, along with the control group, also exhibited changes in the neural processing within the frontoparietal monitoring network. Specifically, following training and practice, older adults exhibited increased activity in inferior parietal regions associated with monitoring load and decreases in medial frontal and inferior parietal regions related to lure success. Though training and practice was met with both increases and decreases in neural activation, the pattern of these findings is consistent with past research on monitoring and false memories. Specifically, retrieval-related monitoring is a neural process that has been consistently found to be supported by frontoparietal regions (Gallo et al., 2006; Quamme et al., 2010). Specifically, parietal regions are thought to guide attentional processes when presented with new information and frontal cortices are thought to resolve incongruent information present in conflicting memory traces (Devitt & Schacter, 2016; Koen & Rugg, 2019; Kurkela & Dennis, 2016; Spaniol & Bayen, 2005). With respect to false memory work, previous studies show that while unrelated lures can be rejected based upon category-level information, the correct rejection of related lures requires greater monitoring and evaluation of details because such lures differ from targets only in terms of specific details that are associated with individual exemplars drawn from the same categories (e.g., dogs, chairs) (Bowman & Dennis, 2015; Coane et al., 2007; Devitt & Schacter, 2016; Schacter & Slotnick, 2004). Thus, the correct rejection of related lures is posited to rely on the detection of highly specific perceptual and/or semantic differences between targets and lures during memory retrieval through memory monitoring processes. Retrieval related monitoring of this type has been shown to rely on neural activity in regions within the frontoparietal retrieval monitoring network (for reviews see Cabeza et al., 2012; Ciaramelli et al., 2008; Wagner et al., 2005). With respect to false memory studies, activation within this frontoparietal network has been found during memory retrieval generally, with increased activation shown for related lures irrespective of the source of the false memory (i.e., semantic, perceptual, source error) (Kim & Cabeza, 2007; Kubota et al., 2006; Kurkela & Dennis, 2016; Okado & Stark, 2003; Schacter et al., 1996; Schacter & Slotnick, 2004; Stephan-Otto et al., 2017; Turney & Dennis, 2017; Webb et al., 2016). The current results continue to extend this prior work to the domain of aging, showing that frontoparietal regions are critical for evaluating related lures throughout the adult lifespan (Bowman et al., 2019; Bowman & Dennis, 2015).

The current results also replicate and extend recent work suggesting that neural patterns within portions of the frontoparietal network maintain discriminable information between targets and perceptual lures (Bowman et al., 2019; Ciaramelli et al., 2008; Gallo et al., 2006). Specifically, at pre-training, patterns of neural activity in the inferior parietal and medial superior frontal gyri reliably discriminated between targets and related lures across the entirety of our sample. While the focus of past work has emphasized processing in visual cortices, the current analyses focused more specifically on neural processing in higher order evaluation and monitoring regions. In doing so, the finding of discriminability within medial superior frontal gyrus and bilateral parietal cortices extends previous work in the field of aging to demonstrate that regions outside of visual cortex can detect differences between old and new information even when there is considerable overlap in the perceptual and semantic properties of the stimuli (see also Lee et al., 2019 for evidence that parietal cortex maintains signals that can reliably discriminate between false memories and correct rejections of related lures). Such evidence suggests that even as information related to item history becomes more semanticized in higher order processing regions, such as frontal and parietal cortices, neural patterns are still discriminable even into older adulthood. These results further support the notion that older adults rely on not just visual information when discriminating between old versus new items, but that they also rely on semantic labels, contextual information, and attentional processes when presented with information of varying history (Ciaramelli et al., 2008; Dehon & Brédart, 2004; Kirchhoff et al., 2012b; Park et al., 1984).

Critically, the current results show not just the involvement of these regions in supporting memory success in aging, but also the ability of older adults to modulate and enhance neural processing within these regions following memory practice and training. While frontoparietal activity has consistently been found in aging studies, age-related deficits within this network have frequently been observed in both general memory (Dulas & Duarte, 2011; Fandakova et al., 2014; Luo & Craik, 2009; McDonough et al., 2013; Mitchell et al., 2013; Velanova et al., 2007) and false memory studies (Bowman & Dennis, 2015; Dennis et al., 2014; Fandakova et al., 2014, 2018), with most studies attributing this finding to age-deficits in monitoring-related memory processes (e.g., Mitchell & Johnson, 2009). For example, Fandakova et al., 2018 found that young, but not older adults modulated activity across cingulo-opercular regions when making false alarms and low-quality correct rejections, consistent with the area’s role in postretrieval monitoring. Additionally, this same research group found that older adults who were able to bring online a more “youth-like” neural profile in regions including middle frontal gyrus, and portions of parietal cortex, were better able to accurately discriminate between targets and related lures (Fandakova et al., 2014). While this and previous work has identified individual differences with respect to the role of frontoparietal cortices in false memory errors (Dennis et al., 2014; Dennis & Turney, 2018; Fandakova et al., 2014; Webb & Dennis, 2019), the current results build upon this earlier work showing the flexibility of this network in older adults. As such, the current finding that recruitment of, and processing within frontoparietal regions, can be modified with training and practice is an exciting step in identifying mechanisms by which age-related memory deficits can be ameliorated.

Interestingly, the foregoing modulation of univariate activity following training came in the form of both increases and decreases in overall activation within the frontoparietal network. While, at first glance, this may appear counterintuitive, the differences across regions align with prior studies observing decreased BOLD amplitude associated with improved, or increased, efficiency of cognitive processes (Lustig et al., 2009). That is, increases were observed in precuneus and angular gyrus related to monitoring difficulty. Within memory retrieval and monitoring, parietal cortices, and specifically the precuneus, have been shown to be associated with episodic source retrieval (e.g., Bonnì et al., 2015; Cabeza & Nyberg, 2000; Lundstrom et al., 2003; Lundstrom et al., 2005) and directing attention to visual input (Cavanna & Trimble, 2006; Trimble & Cavanna, 2008). Additionally, activity within the angular gyrus has been associated with the ability to bring online internal representations that support accurate source retrieval (Ramanan et al., 2018; Rugg & King, 2018; Tibon et al., 2019). As such, increased activity within these regions likely reflects the increased activation of internal representations of encoding memory traces that allow the individual to accurately identify the related lure in a “recall to reject” manner. On the other hand, the contrast of monitoring success was met with decreases in superior medial frontal gyrus, middle frontal gyrus and inferior parietal cortices. Critical to this finding, the superior medial frontal gyrus and inferior parietal cortex have both been consistently linked to false memory errors across a wide variety of tasks (for a meta-analysis see Kurkela & Dennis, 2016), with increased activation observed for false alarms compared to correct rejections. The observed *decreases* in this region following training and practice support participants’ ability to accurately detect the lure (and hence, not false alarm). Thus, the current results extend this prior research, showing that when memory improves and correct rejections increase, there is a decrease in brain activation within regions that typically underlie false memory responses.

The current findings also offer initial evidence that, like univariate activation, participants also receive benefits to indices of neural discriminability. Specifically, regions in parietal, temporal and frontal cortex exhibited relations between baseline classification accuracy and change in classification accuracy. Of particular interest was the finding that time-related improvements in discriminability were negatively related to baseline levels of discriminability. Specifically, individuals that exhibited the lowest overall target-lure discriminability at baseline scanning exhibited the greatest improvements following practice and training. This finding is in line with a breadth of cognitive training work that observes individuals who exhibit the lowest behavioral indices at baseline assessment often benefit the most from cognitive training (Rohegar et al., 2020; Roheger et al., 2020; Schiff et al., 2021; Shaw & Hosseini, 2020; Strobach & Karbach, 2021). Specifically, multiple previous reports have observed a negative correlation between baseline cognitive performance and training-related gains across a number of cognitive domains including attention and episodic memory. It may be that older adults who are at a more optimal level of neural discriminability between targets and lures have less need for improvement. Such results add support to the compensation hypothesis of training-induced improvements, in which older adults who are most at risk of cognitive declines stand to benefit the most from targeted cognitive interventions (Lövdén et al., 2020). Similar results have also been observed in attentional neural processes in participants recovering from traumatic brain injuries (Arnemann et al., 2015).

The current study builds upon other cognitive training of memory experiments examining the neural mechanisms associated with memory improvements in healthy older adults. For example, a recent meta-analysis observed increased BOLD amplitude in parietal cortices in older participants following training (Duda & Sweet, 2020). In more targeted work, Kirchhoff and colleagues (2012), observed increased neural recruitment in medial and dorsolateral portions of frontal cortices associated with improvements in veridical recollection in aging. The authors interpreted this differential activity following training as evidence of the malleability of neural processes driven by behavioral modifications. In the current study, we demonstrate that frontoparietal regions are not only malleable to modifications aimed at increasing veridical memory performance, but also those targeting the reduction of erroneous memory commissions in healthy aging. Likewise, past large-scale cognitive training interventions have exhibited robust effects in improving true memory performance in older adults, in some cases with benefits lasting at least five years (Ross et al., 2016, 2018; Sprague et al., 2019). In the current study we demonstrate that not only are cognitive processes associated with true memory performance modifiable in healthy aging, but also cognitive processes related to false memory performance. Future studies should aim to replicate the observed training-related reduction of false memories in larger and more diverse samples, and at longer intervals (years rather than days), to assess the large-scale efficacy of cognitive training on false memory performance in healthy aging.

Altogether, our findings reflect benefits of the RBMS training, along with practice effects in the control group, as depicted by overall increases and decreases in BOLD amplitude. The absence of a training group x time interaction implies that mere exposure to the false memory paradigm during practice may have been enough to alter neural recruitment in the control group (See supplemental table 1). Furthermore, participants in the control group may have also initiated some type of monitoring strategy due to this exposure. However, false memory paradigm practice was not as effective as RBMS training for reducing false alarm rates. Overall, the current results provide optimism for older adults who may be experiencing poor memory discriminability and increased false memories, as the current results suggest that both neural malleability and behavioral improvements are possible.

## 5.0 Limitations and future directions

The present study demonstrated that RBMS training can reduce older adults’ false memories to a greater extent than non-adaptive control training. However, there are several limitations to this work that should be considered. Given the novelty of using cognitive training to specifically reduce false memories, future replication is necessary. The observed improvements in neural discriminability across groups suggests that, given practice with a related-object memory task, older adults can exhibit shifts in their neural processing. However, the absence of time by group interactions within the neuroimaging analyses suggests that both practice as well as RMBS training led to the modulation of brain activity and neural discriminability. This may be a result of the smaller sample size in the current study or a result of the type of non-adaptive control procedures used (active versus passive control). While we recruited 50 participants, only 38 saw the task through to the end (19 participants within each group). Future work should make every attempt to increase sample sizes. It was also the case that the memory tests performed by the non-adaptive control group were similar to those performed by the RBMS training group.

Specifically, the control group viewed all of the same stimuli and performed the same memory tasks as the RBMS group, absent training instructions and adaptive task difficulty. Future work should explore the relative contributions of instruction vs. practice to improvements in memory discriminability. Finally, future work should examine whether similar strategy-based training during encoding could also reduce false memories.

## 6.0 Conclusions

In conclusion, the current study aimed to reduce false memories in older adults via a retrieval-based monitoring strategy intervention and to investigate the neural correlates of training-associated behavioral changes. We observed a reduction in false memory rates in the RBMS training group but not in a non-adaptive training control group, thereby demonstrating the efficacy and specificity of the retrieval-based monitoring intervention that was designed to reduce memory errors of commission. Neurally, we observed both increases and decreases in BOLD amplitude associated with general and successful monitoring processes within our training and groups across regions within a frontoparietal network. Participants with lower baseline neural discriminability between target and lure items tended to receive the greatest benefits in neural discriminability due to training and practice. Collectively, our results highlight the importance of examining the impact of cognitive training on false memory in older adults, and demonstrate that changes associated with retrieval training and practice are borne out neurally via both alterations in BOLD amplitude and neural discriminability. As such, the current study stands among the first to modulate false memory behavior and associated neural processing in healthy older adults and can provide a useful resource for investigators or clinicians aiming to develop effective methods for reducing older adults’ memory errors.

## Acknowledgements.

We thank the Penn State Social, Life, & Engineering Sciences Imaging Center (SLEIC) 3T MRI Facility. We also thank Chaleece Sandberg, Bradley Wyble, Charles Geier, Frank Hillary, Holly Richardson, Courtney Gerver, and Christina Webb for their assistance in planning and data collection for the current study.

---

**Supplemental Figure 1.**
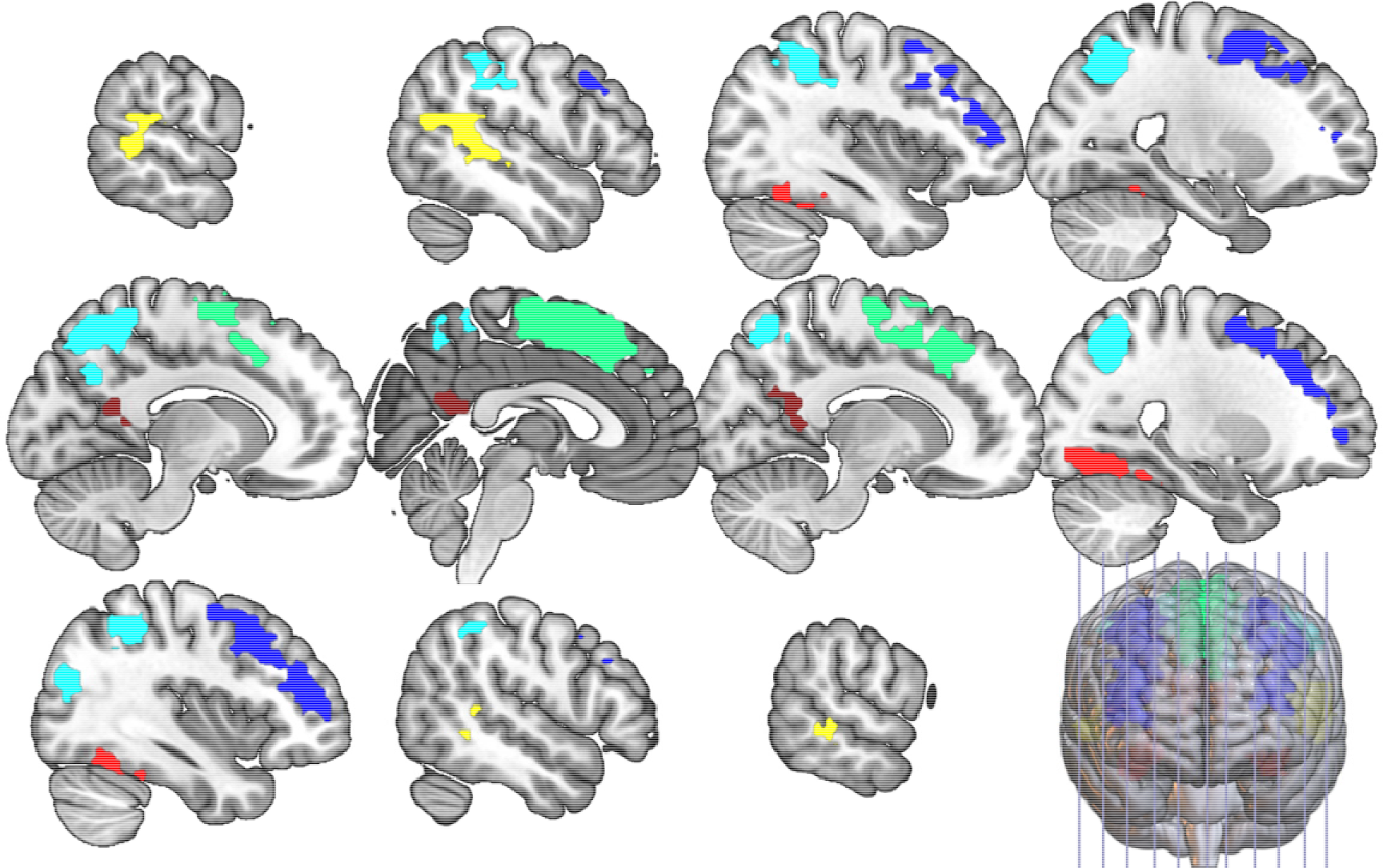
Multivariate classification ROIs. Multivariate ROIs combined from clusters depicting general monitoring and successful monitoring activity at T1. Any overlapping clusters were collapsed to create a single ROI. Yellow = Middle Temporal Gyrus, light blue = Inferior Parietal Gyrus, dark blue = Middle Frontal Gyrus, dark red = Precuneus, light red = Fusiform Gyrus, green = Medial Superior Frontal Gyrus.

**Supplemental Table 1.** The table reports pre-post univariate activity by group for general monitoring (Related Correct Rejections > Unrelated Correct Rejections) and successful monitoring (Related Correct Rejections > Related False Alarms) from T1 to T2. X, y, z represents peak MNI coordinates, k indicates cluster extent, H represents hemisphere, L – left, R-right, t – statistic.

|  | *H* | *MNI Coordinates* |  |  | *k* | *z* |
| --- | --- | --- | --- | --- | --- | --- |
|  |  | *x* | *y* | *z* |  |  |
| *Related Correct Rejection > Unrelated Correct Rejection* |  |  |  |  |  |  |
| Positive Main Effect of Time (Train Only) |  |  |  |  |  |  |
| Precuneus | R | -12 | -70 | 50 | 8 | 3.81 |
| Positive Main Effect of Time (Control Only) |  |  |  |  |  |  |
| No suprathreshold clusters |  |  |  |  |  |  |
| *Related Correct Rejection > Related False Alarm* |  |  |  |  |  |  |
| Negative Main Effect of Time (Train Only) |  |  |  |  |  |  |
| Medial Superior Frontal Gyrus | L | -9 | 14 | 50 | 230 | 4.51 |
| Middle Frontal Gyrus | L | -24 | 11 | 47 | 52 | 4.31 |
| Inferior Parietal Gyrus | R | 48 | -37 | 47 | 30 | 4.18 |
| Precentral Gyrus | R | 27 | -4 | 53 | 20 | 4.03 |
| Inferior Parietal Gyrus | L | -42 | -37 | 44 | 17 | 3.77 |
| Middle Frontal Gyrus | L | -39 | 5 | 53 | 5 | 3.72 |
| Inferior Parietal Gyrus | L | -30 | -49 | 50 | 38 | 3.67 |
| Middle Frontal Gyrus | R | 24 | 14 | 50 | 6 | 3.65 |
| Middle Frontal Gyrus | R | 36 | 5 | 38 | 7 | 3.62 |
| Middle Frontal Gyrus | R | 27 | 8 | 56 | 4 | 3.46 |
| Middle Frontal Gyrus | R | 36 | 17 | 44 | 9 | 3.35 |
| Inferior Frontal Gyrus | L | -45 | 23 | 32 | 8 | 3.34 |
| Negative Main Effect of Time (Control Only) |  |  |  |  |  |  |
| Inferior Parietal Gyrus | L | -57 | -31 | 44 | 4 | 3.42 |

