## SupplementalFigureTable for "Investigation of the neural effects of memory training to reduce false memories in older adults: Univariate and multivariate analyses"

**Supplemental Tables and Figures**

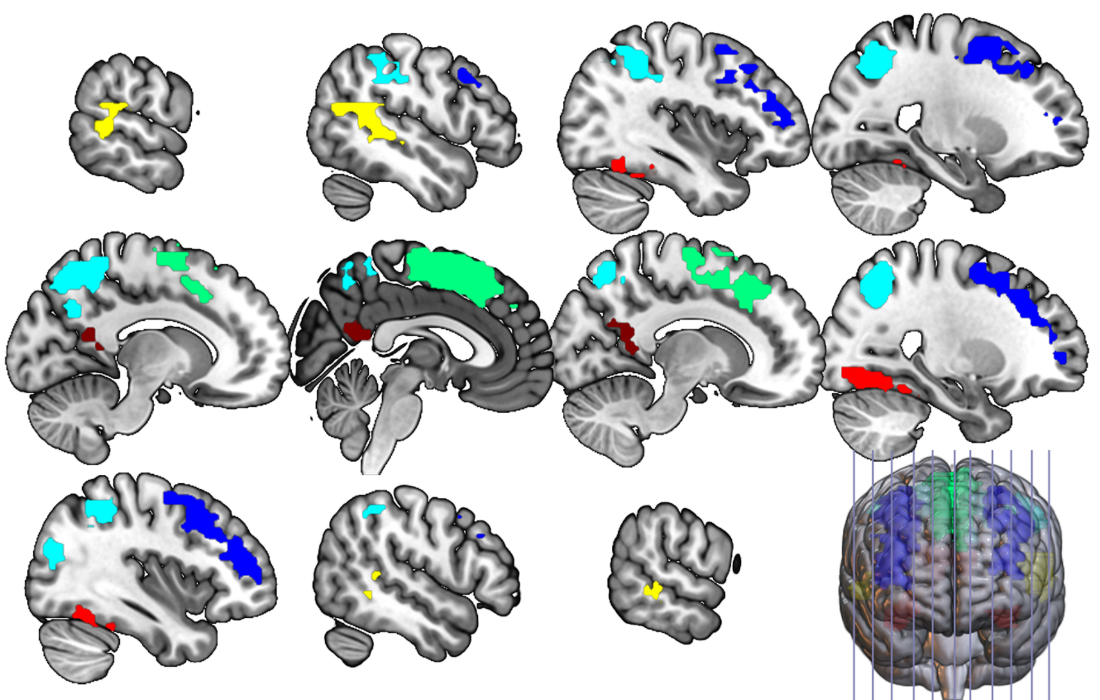

**Supplemental Figure 1.** Multivariate classification ROIs. Multivariate ROIs combined from clusters depicting general monitoring and successful monitoring activity at T1. Any overlapping clusters were collapsed to create a single ROI. Yellow = Middle Temporal Gyrus, light blue = Inferior Parietal Gyrus, dark blue = Middle Frontal Gyrus, dark red = Precuneus, light red = Fusiform Gyrus, green = Medial Superior Frontal Gyrus.

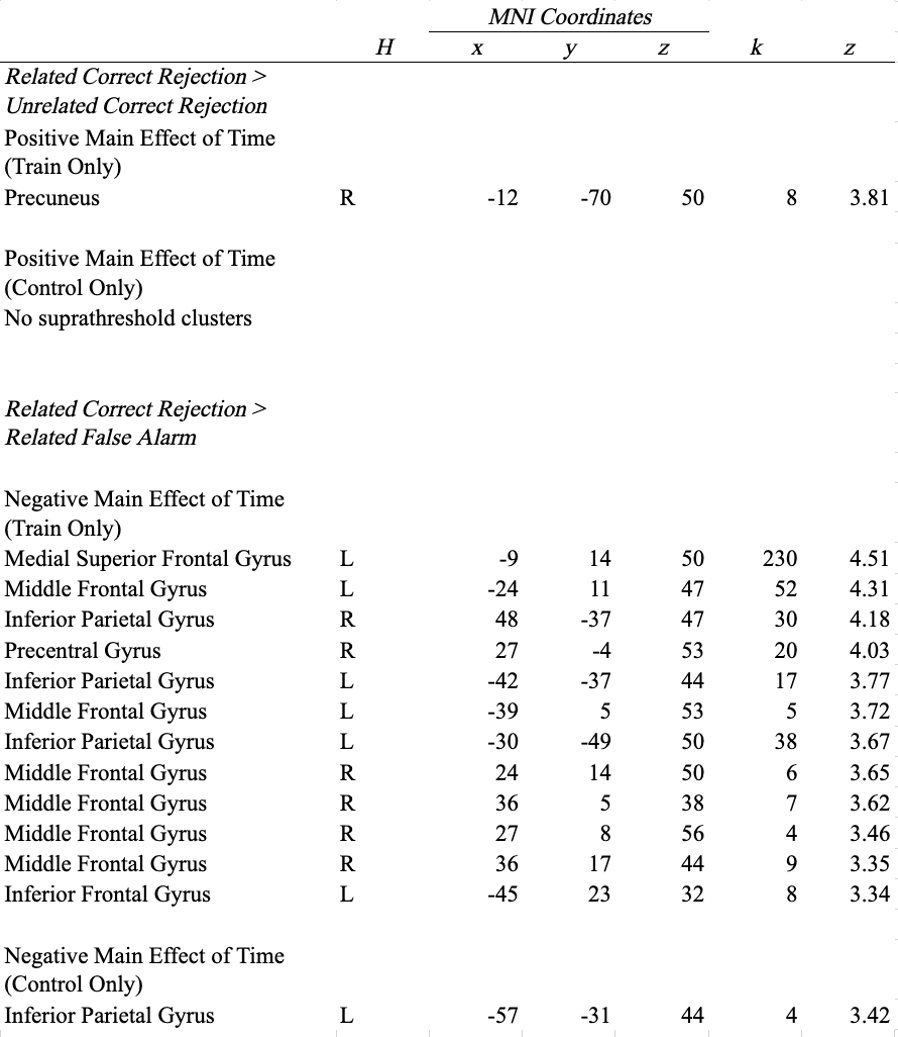

**Supplemental Table 1.** The table reports pre-post univariate activity by group for general monitoring (Related Correct Rejections > Unrelated Correct Rejections) and successful monitoring (Related Correct Rejections > Related False Alarms) from T1 to T2. X, y, z represents peak MNI coordinates, k indicates cluster extent, H represents hemisphere, L – left, R- right, t – statistic.
